# Metazoan ribotoxin genes acquired by Horizontal Gene Transfer

**DOI:** 10.1101/071340

**Authors:** Walter J. Lapadula, Paula L. Marcet, María L. Mascotti, María V. Sánchez Puerta, Maximiliano Juri Ayub

## Abstract

Ribosome inactivating proteins (RIPs) are RNA *N*-glycosidases that depurinate a specific adenine residue in the conserved sarcin/ricin loop of 28S rRNA. These enzymes are widely distributed among plants and their presence has also been confirmed in several bacterial species. Recently, we reported for the first time *in silico* evidence of RIP encoding genes in metazoans, in two closely related species of insects: *Aedes aegypti* and *Culex quinquefasciatus*. Here, we have experimentally confirmed the presence of these genes in mosquitoes and attempted to unveil their evolutionary history. A detailed study was conducted, including evaluation of taxonomic distribution, phylogenetic inferences and microsynteny analyses, indicating that the culicine RIP genes derived from a single Horizontal Gene Transfer (HGT) event, probably from a Cyanobacterial donor species. Moreover, evolutionary analyses show that, after transference, these genes evolved under purifying selection, strongly suggesting that they play functional roles in these organisms. In this work we confirm the presence of RIP genes in Culicinae species, and show solid evidence supporting the hypothesis that these genes are derived from a single prokaryotic transferred gene through HGT. In addition, clear evidence of purifying selection pressure has been recorded, supporting the hypothesis that these genes are functional within this subfamily.

## Introduction

Ribosome inactivating proteins (RIPs, EC 3.2.2.22) irreversibly modify ribosomes through the depurination of an adenine residue in the conserved alpha-sarcin/ ricin loop of 28S rRNA (1-4). This modification prevents the binding of elongation factor 2 to the ribosome, arresting protein synthesis (5, 6). The occurrence of RIP genes has been experimentally confirmed in a wide range of plant taxa, as well as in several species of gram positive and negative bacteria (7-9). Additionally, the exponential increase of information in databases has suggested the presence of these toxin genes in Fungi, Cyanobacteria and Metazoan lineages (10-12). Although several RIPs have been extensively studied at the biochemical level, their biological roles remain open to speculation. In some cases, it seems reasonable to predict their functions. For instance, the high toxicity of ricin supports an antifeedant role, whereas shiga and shiga-like toxins are strong virulence factors for their harboring bacteria. Antiviral and other defense activities have been postulated for other plant RIPs, but no concluding evidence has been obtained. Recently, the RIP of the symbiotic *Spiroplama* in *Drosophila neotestacea* was shown playing a defensive role in preventing a virulent nematode from infecting this insect (13).

In a previous work, we have described that phylogeny of RIP genes show incongruence with those of species. Most of these inconsistencies can be explained by gene duplication, loss and/or lineage sorting (11). Another mechanism leading to phylogeny incongruence is horizontal gene transfer (HGT); namely the non-genealogical transmission of DNA among organisms. HGT is accepted as an important force driving prokaryotic genome evolution (14, 15). In contrast, its impact on genomes from multicellular eukaryotes, in particular animals, is largely controversial (16, 17). To be maintained permanently in animal species, heritable changes (*i.e.* the transferred gene) must be incorporated into germline cells and transmitted to the offspring. Nevertheless, in the particular case of herbivore arthropods, HGT has been postulated to play a role in the adaptation to phytophagy, including the efficient assimilation and detoxification of plant produced metabolites (18).

Detection of *bona fide* HGT derived genes is not trivial, and careful data revision is required for its corroboration. Many cases of putative foreign genes have been shown, after further revision, to result from artifacts or misinterpretations, such as contamination of genomic data, incomplete sampling of sequences and/or taxa, incorrect phylogenetic inferences and hidden paralogy. Two emblematic cases illustrating these issues are the initial conclusion that the human genome contained a high percent of bacterial derived genes (19), and the recent claim that tardigrade genomes contain significant amounts of foreign DNA (20). In both cases, subsequent sounder analyses demonstrated that contamination or incomplete sampling better explained the available data (21, 22). Consequently, careful case-by-case analyses of HGT candidates are required for their efficient detection. To do so, independent evidences and alternative evolutionary scenarios should be taken into account.

Based on the previous finding of *in silico* evidence of the presence of RIP genes in two closely related species of mosquitoes, we aim to confirm the presence and determine the location of RIP genes in species of the Culicinae subfamily. Moreover, we show solid evidence supporting the hypothesis that these genes derive from a single prokaryotic transferred gene.

## Materials and Methods

### PCR experiments

PCR experiments were conducted to confirm the presence of RIP encoding sequences in selected organisms. Individuals of *C. quinquefasciatus* strain JHB were obtained from the MR4 colony (Malaria Research and Reference Reagent Resource Center) of the Center for Diseases Control and Prevention (CDC). Genomic DNA from these specimens was extracted employing the protocol previously described by Collins (23). Genomic DNA from wild specimens of *C. molestus, C. pipiens* and *C. torrentium* was kindly provided by Dr. Stefanie Becker (Bernhard Nocht Institute for Tropical Medicine, Hamburg) (24). Primer sets were designed to amplify the full-length RIP encoding sequence of *C. quinquefasciatus* and also to identify the predicted neighbor gene (S1 Table). High fidelity *Phusion* DNA polymerase (New England Biolabs) was used. PCR conditions were: initial denaturation 30 s at 98°C, followed by 35 cycles of denaturation (10s at 98°C), annealing (30s at Ta°C see S1 Table) and extension (30s at 72°C), final extension of 10 min at 72°C. PCR products were cloned into the pGEMT-easy vector (Promega) following standard methods, and sequenced. The obtained sequences are available with the following GenBank codes: KX674699, KX674697, KX644696 and KX674698.

### Multiple sequence Alignments (MSAs) and phylogenetic inferences

We used the amino acids sequences of previously reported RIPs (11) as probes for iterative BLAST searches. All collected sequences were employed to construct MSAs using MAFFT 7 on-line server (http://mafft.cbrc.jp/alignment/server/). MSAs were manually edited to remove gaps. Phylogenetic trees were obtained employing Bayesian inference using Mr. Bayes 3.2 software (25). A mixed amino acid substitution model was set up, 4 gamma categories and a proportion of invariable sites were considered. The analyses were concluded after 1,000,000 generations when the split frequency was < 0.02. FigTree 1.4.2 software was used to visualize and edit the obtained trees.

### Genomic context analyses

The whole contigs containing RIPs of *C. quinquefasciatus* (DS232037), *Aedes aegypti* (NW001810221) and *Anopheles gambiae* (chromosome 3L) were subjected to BLASTx searches on different protein databases to identify individual genes upstream and downstream from the RIP open reading frame (ORF). The retrieved sequences were subjected to reciprocal BLASTp and tBLASTn searches in order to determine putative orthologous sequences. The obtained orthologs were confirmed using the comparative genomic tool available at VectorBase website (https://www.vectorbase.org/).

### Evolutionary analyses

Nucleotide encoding RIP sequences found in mosquitoes were aligned by codons, using PAL2NAL algorithm (26). Then, the synonymous and non-synonymous substitution rates were estimated employing SLAC, FEL and REL tests available in the Datamonkey Package (27, 28). Finally sequence logo diagram was constructed employing the codon alignment using the Weblogo software (29) and codons under purifying selection were highlighted.

## Results

### *Culex* spp genomes harbor RIP encoding genes

Recently, we have found *in silico* evidence for the presence of RIP genes in two closely related species of metazoans: *Aedes aegypti* and *Culex quinquefasciatus* (11). These intriguing findings led us to design experimental strategies to confirm their presence by ruling out possible database artifacts (*i.e*. contamination). For this purpose, genomic DNA was obtained from a pool of four mosquitoes of *C. quinquefasciatus* strain JHB. Then, two independent PCR experiments were designed to demonstrate the presence of the RIP gene, and to confirm its physical linkage to the predicted neighbor gene, which is an intron-containing metazoan-derived gene (XM_001850822). Fig 1 shows that both PCR products presented the expected size. Also, further cleavage of both amplicons with EcoRI yielded the predicted patterns, confirming their identity.

**Fig 1:**
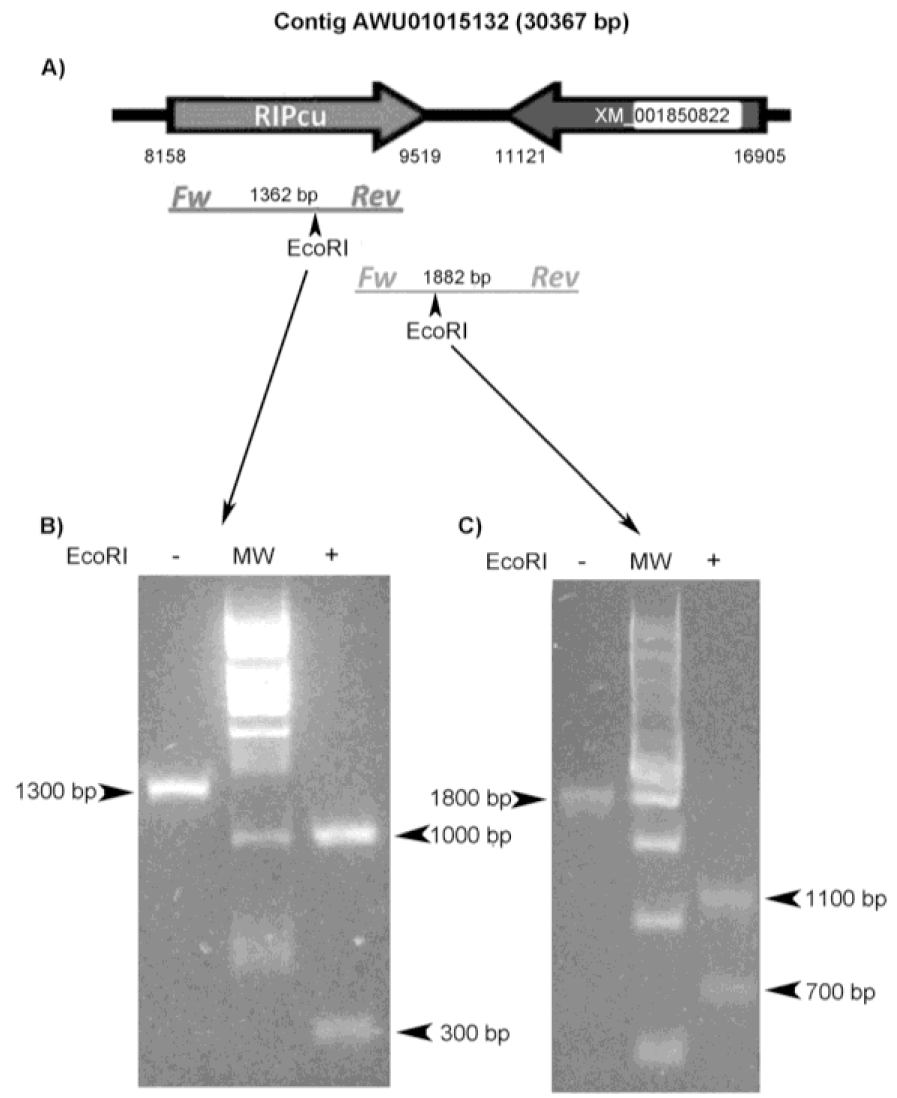
Experimental confirmation of the presence and location of RIP gene in *C. quinquefasciatus* JHB genome. **(A)** Schematic representation of a fragment of the contig AAWU01015132 depicting the RIP gene (RIPcu) and its closest neighbor gene (XM_001850822). Expected amplicons and relevant EcoRI restriction sites are also depicted. **(B)** RIP ORF was amplified by PCR and the product was analyzed by gel electrophoresis before (lane 1) and after EcoRI treatment (lane 2). **(C)** A genomic fragment of 1,882 bp linking the RIP gene with its neighbor gene was amplified and electrophoresed before (lane 1) and after EcoRI treatment (lane 2).

Once the presence and location of the RIP sequence in *C*. *quinquefasciatus* genome were experimentally confirmed, we successfully amplified the full length RIP coding sequences (∼1300 bp) of the closely related species *C. pipiens*, *C. molestus,* and *C. torrentium* (S1 Fig). PCR products were cloned and sequenced, and the obtained sequences were aligned. As expected, nucleotide sequences showed high similarity (93-97% of identity at nucleotides and amino acids levels) in relation to the reported sequence in the *C. quinquefasciatus* genome database (S2 Fig). The sequence obtained from the *C. quinquefasciatus* JHB MR4-CDC repository revealed an in-frame, three-nucleotide (ACC) insertion (encoding an additional Thr residue). Interestingly, this sequence also harbors a ten-nucleotide frame-shifting deletion (nt 542-551) generating a premature stop codon (S2 Fig). By direct sequencing of PCR products from six individual specimens of *C. quinquefasciatus* JHB from the MR4-CDC colony, we confirmed that all these individuals were homozygous for the deletion, strongly suggesting that this null mutation was fixed in this colony (S3 Fig).

### Culicine RIP genes are monophyletic and syntenic

We have previously shown that RIP genes from particular lineages (*e.g.* monocots or dicots, bacteria and fungi) are not monophyletic. Moreover, many RIP clades include sequences belonging to largely distant taxa (11, 30). Based on this evidence, we postulated that the evolutionary history of RIP genes is consistent with the existence of several ancient paralogues, followed by multiple lineage-specific gene duplications and losses (11). However, metazoan RIP genes are particularly interesting because they are restricted to closely related insects of the subfamily Culicinae. As can be seen in Fig 2, RIPs from *Culex quinquefasciatus* and *Aedes aegypti* form a well supported clade [posterior probability (PP): 1]. Monophyly of culicine RIPs, along with their apparent narrow taxonomic distribution, suggest that these genes are derived from a rather recent, single ancestral sequence. In order to test this, microsynteny analyses were carried out using scaffolds from *Aedes aegypti, Culex quinquefasciatus* and *Anopheles gambiae* (the closest relative specie lacking RIP genes). As expected, partially conserved synteny blocks were identified in the three species (Fig 3 and S2 Table).

**Fig 2:**
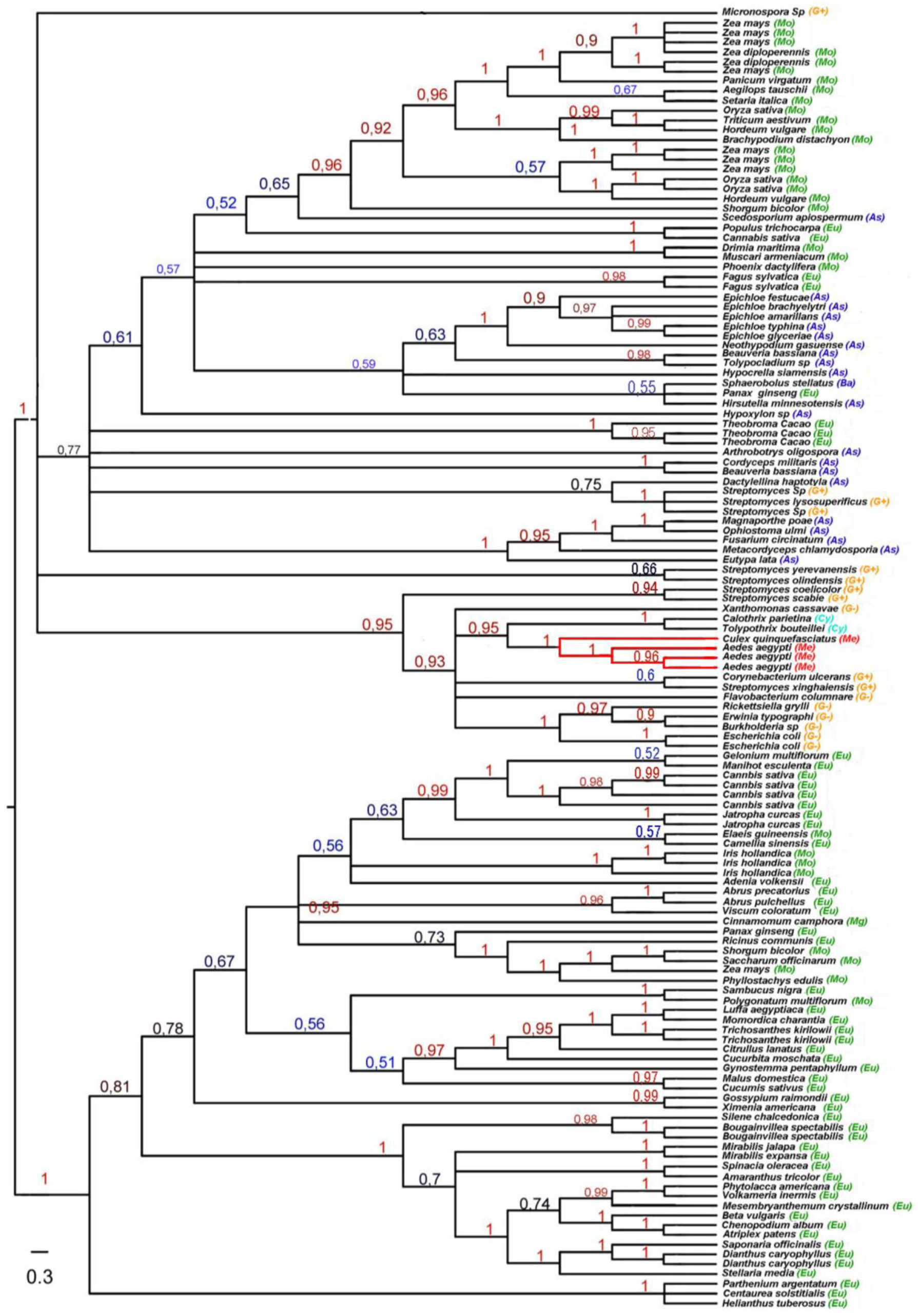
Phylogenetic tree of RIPs protein family. The tree was constructed employing Bayesian Inference. Numbers above the branches indicate the posterior probabilities (PP) values. Names of species are indicated for each sequence in the tree. Lineages are indicated by different color codes as follows: green (Plant), blue (Fungi), red (Metazoan), orange (Bacteria) and magenta (Cyanobacteria). The clade of dipterous RIPs is emphasized with red branches.

**Fig 3:**
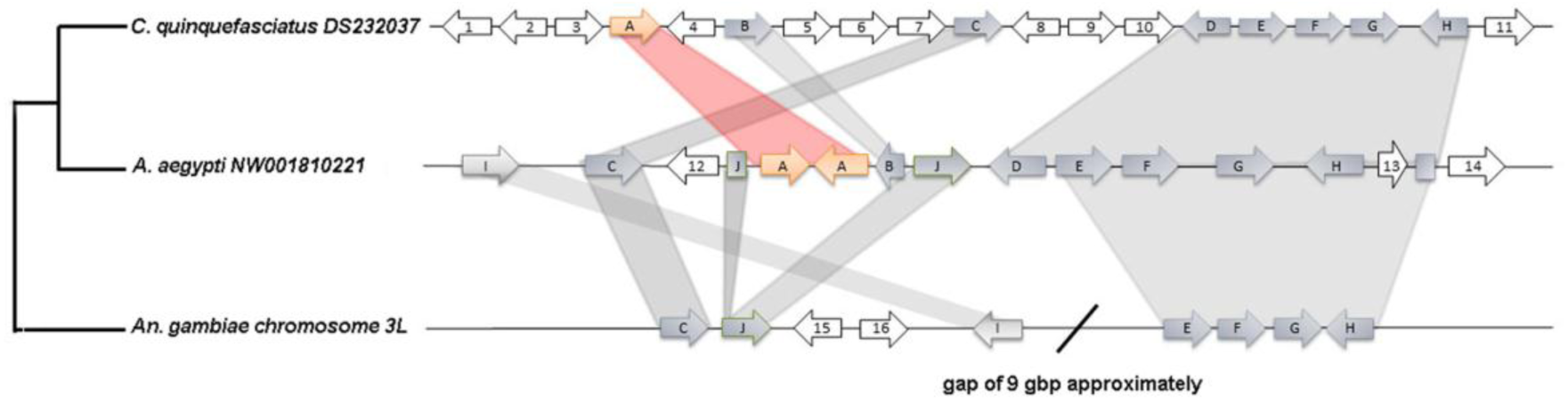
Schematic representation of the genomic context between *Culex quinquefasciatus* (DS232037), *Aedes aegypti* (NW001810221) and *Anopheles gambiae* (chromosome 3L). Greys shadows link conserved syntenic ORFs. RIP genes are represented with orange arrows. Additional information about each gene is available in Table S2.

### Culicine RIP genes are derived from a unique HGT event

Recent sequence homology searches further confirmed that metazoan RIP genes are restricted to the Culicinae subfamily. In addition to the above described genes, we found *in silico* seven RIP genes in *Aedes albopictus* (GenBank: KXJ78156, KXJ78155, KXJ78158, KXJ73132, KXJ78157, KXJ73133 and KXJ73764), which are clearly syntenic with *Aedes aegypti* RIPs (Data File S1) and two transcriptomic sequences from another Culicine mosquito; *Armigeres subalbatus*, partially covering the ORFs (GenBank: EU212208, EU211398). In light of these findings, two alternative hypotheses were postulated:

i. These genes have been vertically inherited from the metazoan cenancestor, and were purged from other metazoan genomes by a number of independent gene loss events.
ii. These genes are derived from a unique HGT event which took place in the common ancestor of the *Culex* and *Aedes* species.

The minimal number of independent gene loss events required was determined in order to evaluate the plausibility of the vertical transmission hypothesis (see Data File S2 for details). Following a conservative approach, at least 15 gene losses should be postulated, from Bilateria to Culicidae, to explain the narrow taxonomic distribution of RIP genes in Metazoa. The alternative HGT hypothesis involves a single gene loss event in the lineage to extant metazoans followed by a single recent HGT event in the ancestor of *Culex* and *Aedes*, yielding a more parsimonious evolutionary scenario.

Based on the strong evidence supporting that culicinae RIPs are derived from an HGT event, a search for possible donors was conducted. The phylogeny of RIPs shows that culicine RIPs form a well supported monophyletic group (PP: 1) embedded within bacterial sequences (Fig 2), suggesting a prokaryotic origin of these sequences. The lack of introns in culicine RIPs also supports this idea. Moreover, BLASTp and tBLASTn searches using mosquito RIPs as queries yielded sequences belonging to *Tolypothrix bouteillei* (Cyanobacteria), *Calothrix parietina* (Cyanobacteria) and *Spiroplasma* spp (Tenericutes). In order to perform a more robust study we focused on culicine, including the recently reported sequences from *A. albopictus*, and on bacterial RIPs. Phylogenetic analysis showed that culicine RIPs form a monophyletic group (PP: 0.96) with sequences from *Spiroplasma* spp and Cyanobacteria species (Fig 4). Thus, these organisms –or others closely related- could have been potential donors.

**Fig 4:**
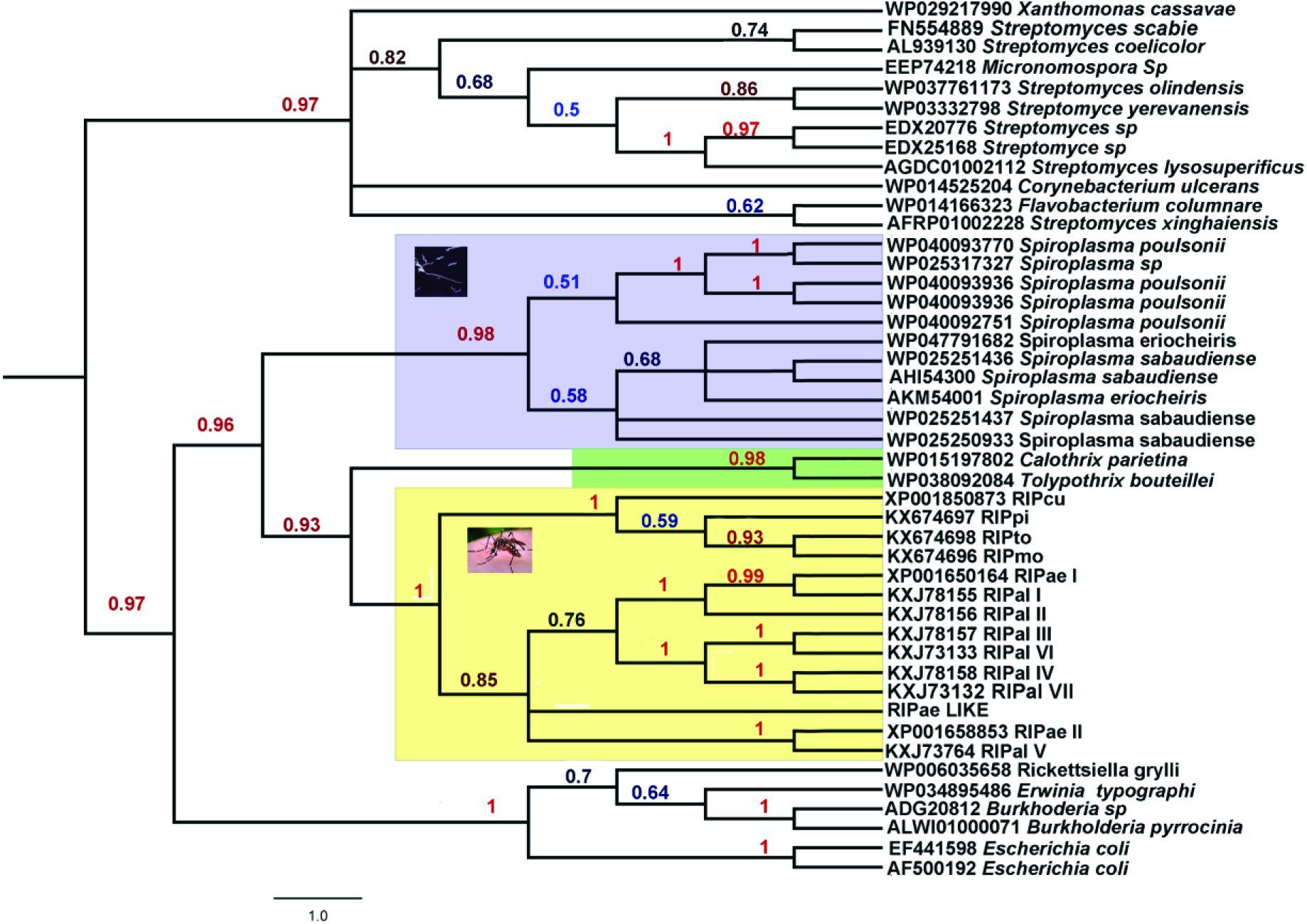
Phylogenetic relationships among metazoan and bacterial RIPs. Numbers above branches indicate Posterior Probabilities (PP) values. Light blue, green and yellow backgrounds indicate *Spiroplasma*, *Cyanobacteria* and mosquitoes RIPs, respectively.

### Horizontally acquired RIP genes evolve under purifying selection pressure

Except for a few well characterized potent toxins (*e.g*. ricin, shiga and shiga-like), the physiological role of most RIPs remains unknown (31). The defensive role demonstrated for *Spiroplasma* RIP genes in *Drosophila neotestacea* (13) invites us to postulate a biological function for HGT acquired RIP genes in mosquitoes. Therefore, we searched for signs of selective pressure on HGT derived sequences as reliable evidence of functionality in insects. To do this, all RIP encoding sequences from *C. quinquefasciatus*, *C. molestus*, *C. pipiens*, *C. torrentium*, *A. aegypti* and *A. albopictus* were aligned (Dataset S1). Interestingly, the observed INDELs were always multiple of three nucleotides, strongly suggesting that frame-shifting mutations were actively purged by selection pressure. Fig 5 shows a Logo representation of the MSA, where a higher conservation degree of first and second bases of each codon seems to be the rule. Moreover, an integrative analysis of synonymous *vs* non-synonymous substitutions employing three different methods (SLAC, FEL and REL) showed purifying (negative) selection for 64 codons, including most of the amino acids forming the active site (Fig 5 and S3 Table).

**Fig 5:**
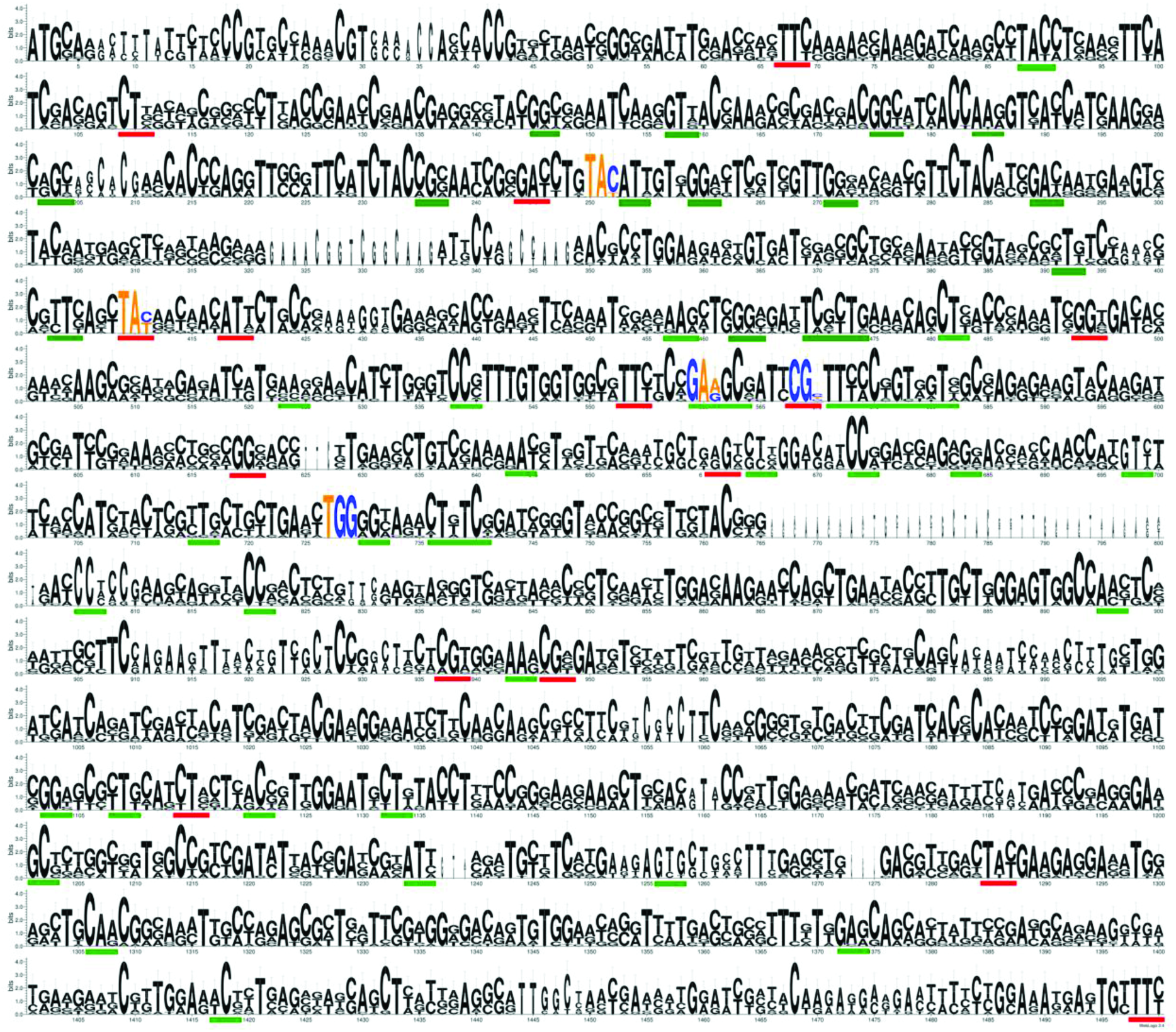
Logo representation of the MSA used to analyze the synonymous *vs* non-synonymous substitution rates. Codons forming the active site are indicated by colored nucleotides (A and T: yellow, G and C: blue). Codons under significant purifying selection determined by the three tests (SLAC, FEL and REL) are underlined in red color. Codons detected to be under significant purifying selection by two out of the three tests, are underlined in green color.

## Discussion

Ribosome inactivating proteins form a very interesting protein family displaying a patchy taxonomic distribution. As it was mentioned above, in a previous report we have found *in silico* evidence of the presence of RIP genes in two closely related species of mosquitos (11). Due to the uniqueness of this finding, in this work we have experimentally confirmed the presence and location of RIP genes in the *C. quinquefasciatus* genome, as well as in other *Culex* species. Moreover, new exhaustive searches by BLAST and HMMER on metazoan databases, revealed the presence of additional homologous genes in *A. albopictus* and *Armigeres sulbabatus*, confirming that the RIP gene family is taxonomically restricted to the Culicinae subfamily (Data File S1 and S2).

Currently, there is not a “gold standard” methodology for automatic and reliable detection of HGT. Moreover, several reports claiming the presence of foreign genes have been undermined after more thorough analyses due to artifacts or misinterpretations (19, 32-34). Therefore, careful integration of information derived from taxonomic distribution, phylogenetic inferences and biological information is needed to detect *bona fide* horizontally acquired genes. The evidence obtained in this work shows that the most plausible origin of culicine RIPs is a single HGT event to the cenancestor of *Culex* and *Aedes* genera. This model is supported by the monophyly of metazoan RIPs and their very narrow taxonomic distribution, the gathering in a clade along with prokaryotic sequences and the shared genomic context among Culicinae species.

According to phylogenetic inferences (Figs 2 and 4), the donor of the RIP gene was, most likely, a prokaryotic organism. An obvious donor candidate for insects is *Wolbachia* spp, since several HGT events between these bacteria and arthropods have been clearly documented (35-37). It is expected that animal genomes are marginally affected by HGT because of the separation of the germline from somatic cells. This barrier to HGT, known as Weismann barrier is not present in the case of bacteria infecting germline cells, as *Wolbacchia* spp, which is consistent with the relatively high number of *Wolbacchia* to insect HGT events (17). However, no RIPs encoding sequences can be found in any of the *Wolbachia* spp databases (including 27 fully sequenced genomes). Interestingly, homology searches using mosquitoes RIPs as probe yielded significant similarity to several *Spiroplasma* genes (Fig 5). The facts that *Spiroplasma* species lack cell walls and that they are frequent endosymbionts of arthropods make them logical donor candidates. However, considering *Spiroplasma* spp as donor involves two major drawbacks. *Spiroplasma* spp coding sequences harbor very low GC content (around 23%), whereas RIP genes from Diptera range from 44.8% to 55.6% (Table 1). Secondly, *Spiroplasma* spp and some species of *Millicutes* use a non-universal UGA tryptophan codon. This variation in the genetic code is presumed to have occurred in the early divergence of these genera (dating 250 mya approximately) (38), and the Culicinae subfamily has diverged more recently (between 51 to 204 mya) (39, 40), the transferred genes containing the non-universal UGA tryptophan codons would be read as a stop. Although CG content of a transferred functional gene could be gradually modified by amelioration, the reversion of several nonsense codons to Trp does not seem plausible. Therefore, these pieces of evidence lead us to reject the hypothesis that Diptera RIP genes are derived from *Spiroplasma*.

**Table 1.**
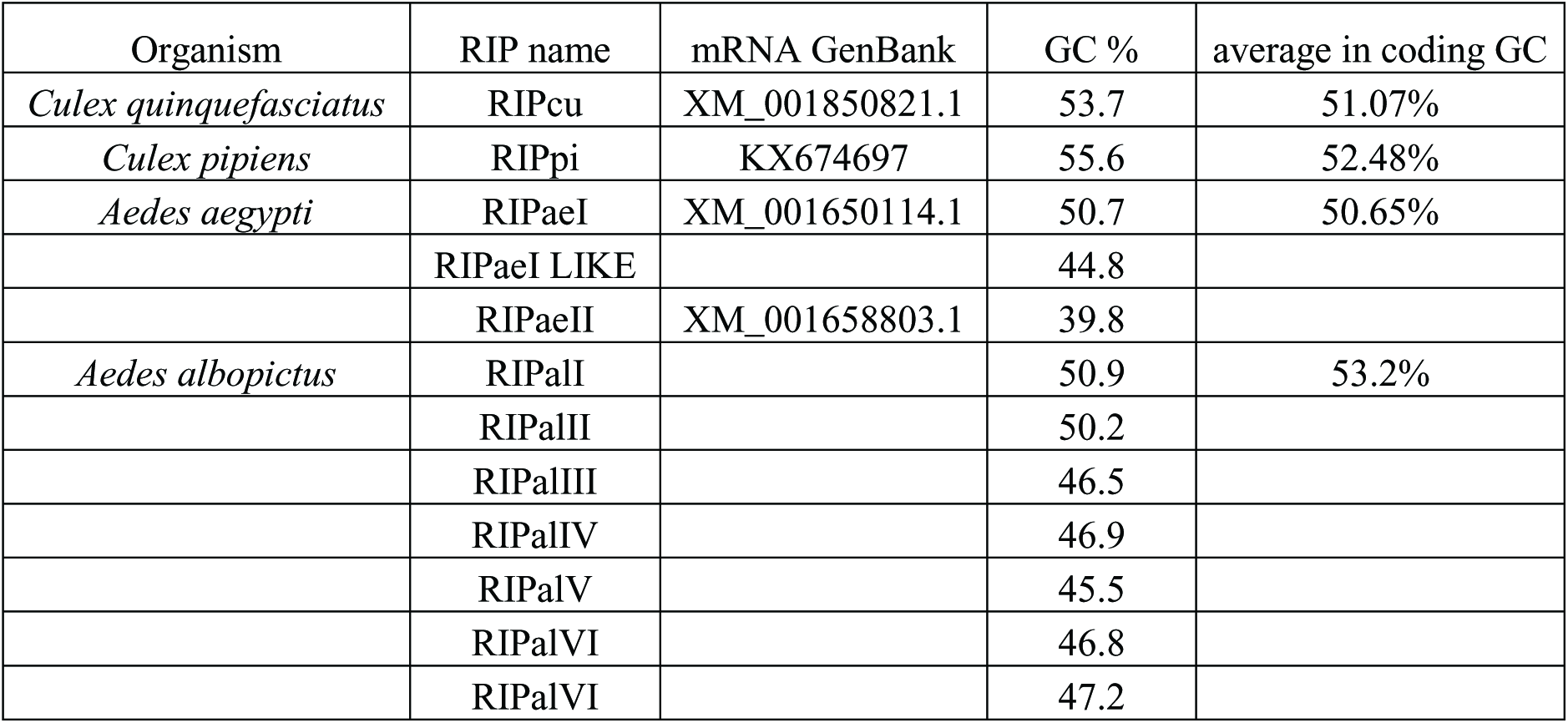
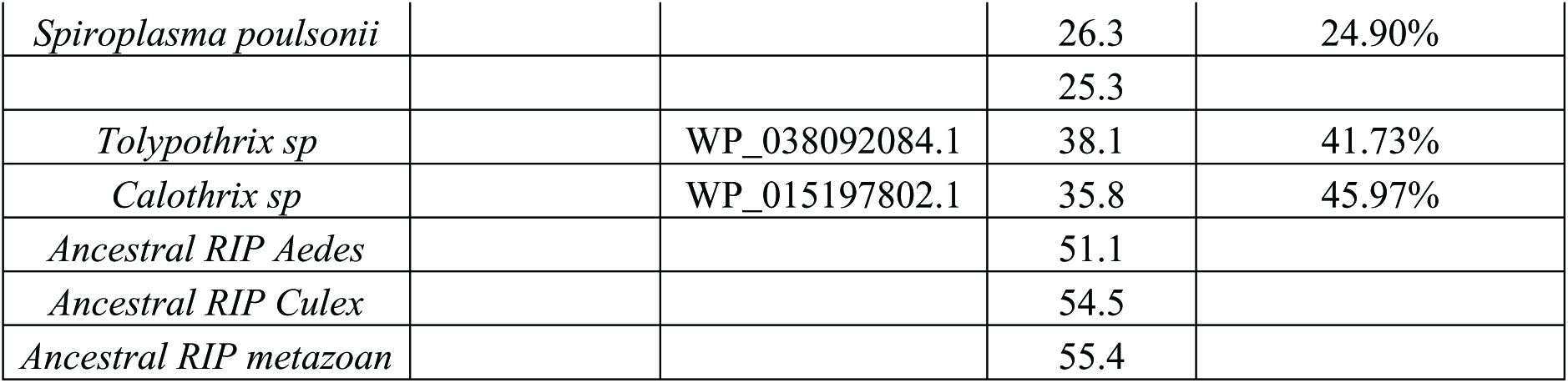
**GC content analysis.** The GC content of RIP genes was calculated using DNA/RNA GC Content Calculator (http://www.endmemo.com/bio/gc.php). The average of GC content in coding region was obtained from the codon usage database (http://www.kazusa.or.jp/codon/).

The phylogenetically closest sequences to culicine RIPs belong to Cyanobacteria species (*Tolypothrix bouteillei* and *Calothrix parietina*). Cyanobacteria constitute a significant fraction of the microbiota at breeding sites of mosquitoes. Remarkably, Cyanobacteria species account for 40% of the bacterial midgut content of larval and pupal stages in *An. gambiae* (41). Moreover, *Calothrix* sp has been detected in the midgut bacterial flora of *An. stephensi* in larval stage (42). In addition, GC content of Cyanobacteria genomes is closer to the Diptera RIPs (Table 1). Altogether; the presented phylogenetic inferences, the shared ecological niches between these bacteria and insects, and the colonization of mosquitoes in their early developmental stages (41) strongly suggest that Cyanobacteria are the most plausible donor species of RIP genes through HGT to the Culicinae subfamily. In line with Huang (17), we postulate that Weisman barrier is, if not absent, markedly weakened in the egg, pupa and larva stages of mosquitoes. Consequently, these early developmental stages of mosquitoes could be particularly prone to the acquisition of heritable foreign genes by environmental bacteria.

An important question about the HGT derived genes is their fate. In other words, the issue at stake is whether “foreign” genes will impact on fitness, and if so, to what extent these genes will be affected by natural selection and/or genetic drift. Probably, mosquitoes RIP genes display a defensive role which has helped them to be fixed by natural selection. However, the fact that individuals from the *C. quinquefasciatus* MR4 colony are homozygous for a null mutation of RIP gene shows that this gene is not essential for viability under laboratory conditions. On the other hand, all mosquitos’ sequences have several codons under purifying selection, which is also reflected by the occurrence of the majority of changes at the third position of codons (Fig 5). According to the nearly-neutral evolutionary theory, slightly deleterious mutations can be fixed in populations with low effective size (as in the case of laboratory colonies). On the contrary, these mutations are efficiently purged from larger populations, such as natural populations (43). This seems to be the case of culicine RIPs, since null mutant *C. quinquefasciatus* are viable in captivity, while clear evidence of selection pressure on coding sequences in wild specimens has been found.

## Conclusions

We have found solid evidence suggesting that culicine RIP genes derive from a single ancestral sequence acquired by HGT from a prokaryotic organism, probably a Cyanobacterium. Moreover, we have also demonstrated that these genes have evolved, after transference, under purifying selection, implying a functional role in the host organism.

## Acknowledgements

The DNA samples of *C. molestus*, *C. pipiens* and *C. torrentium* were kindly gifted by Dr. Stefanie Becker. The authors acknowledge the Malaria Research and Reference Reagent Resource Center (MR4, CDC) for providing the following mosquito strains through BEI Resources, NIAID, NIH: *Culex quinquefasciatus*, strain JHB, NR-43025. The authors are also grateful to Dr. Jimena Juri Ayub for her technical support. We would also like to thank GAECI for their services. The findings and conclusions in this manuscript are those of the authors and do not necessarily represent the views of the Centers for Disease Control and Prevention (CDC).

## Supplementary Information

**S1 Table:**
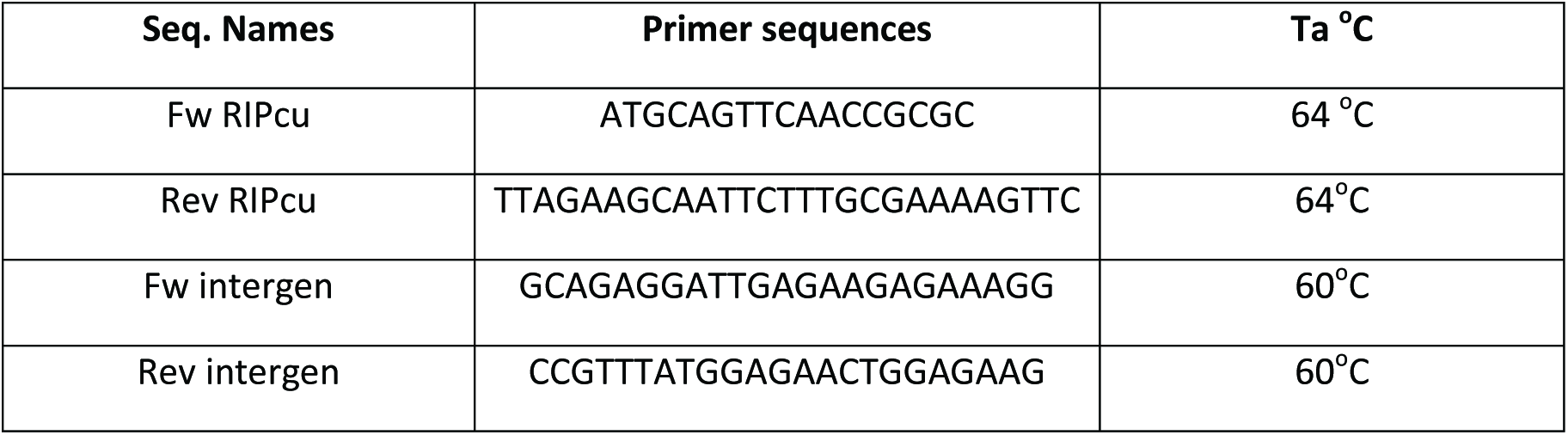
Primer sequences and annealing temperatures (Ta) are indicated in second and third column respectively.

**S1 Fig:**
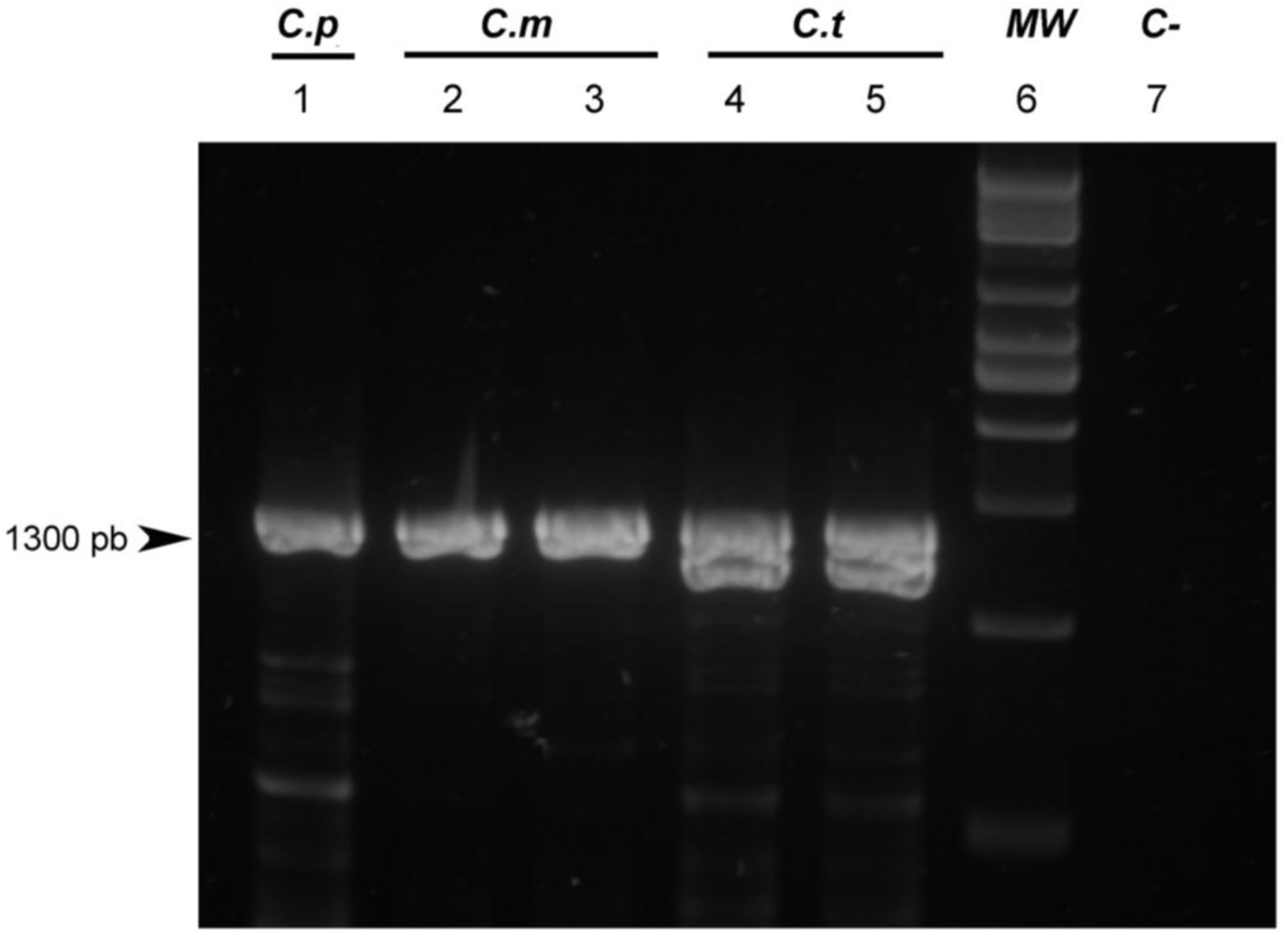
PCR amplification of RIP genes in species belonging to the *Culex* genera. *C. pipiens* (C.p, lane 1), C. *molestus* (C.m, lanes 2 and 3) and *C. torrentium* (C.t, lanes 4 and 5). Lane 6: 1Kb molecular weight size marker, lane 7: negative control.

**S2 Fig:**
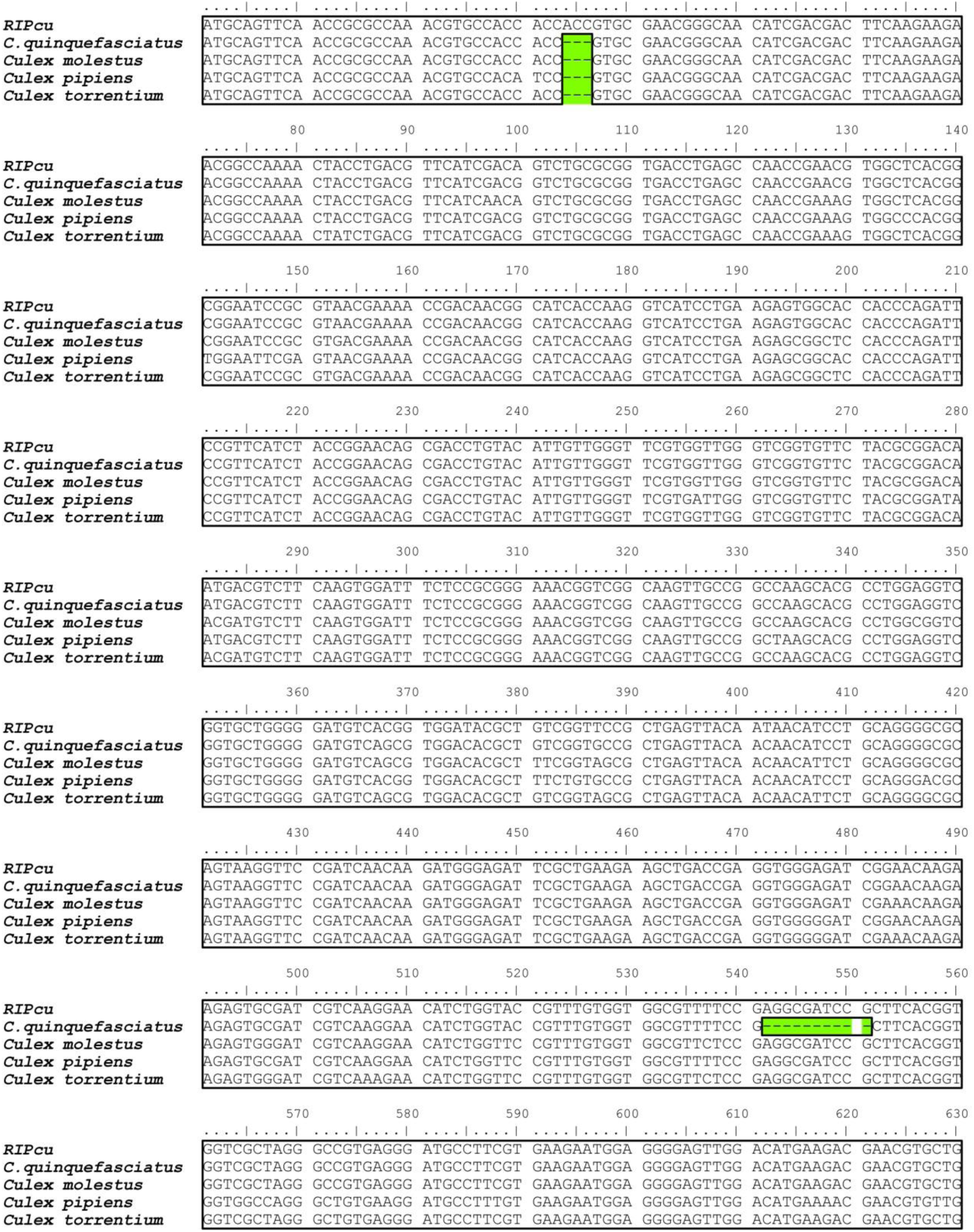
MSA of partial RIPs genes found in *Culex* ssp. Gaps are indicated in green color boxes.

**S3 Fig:**
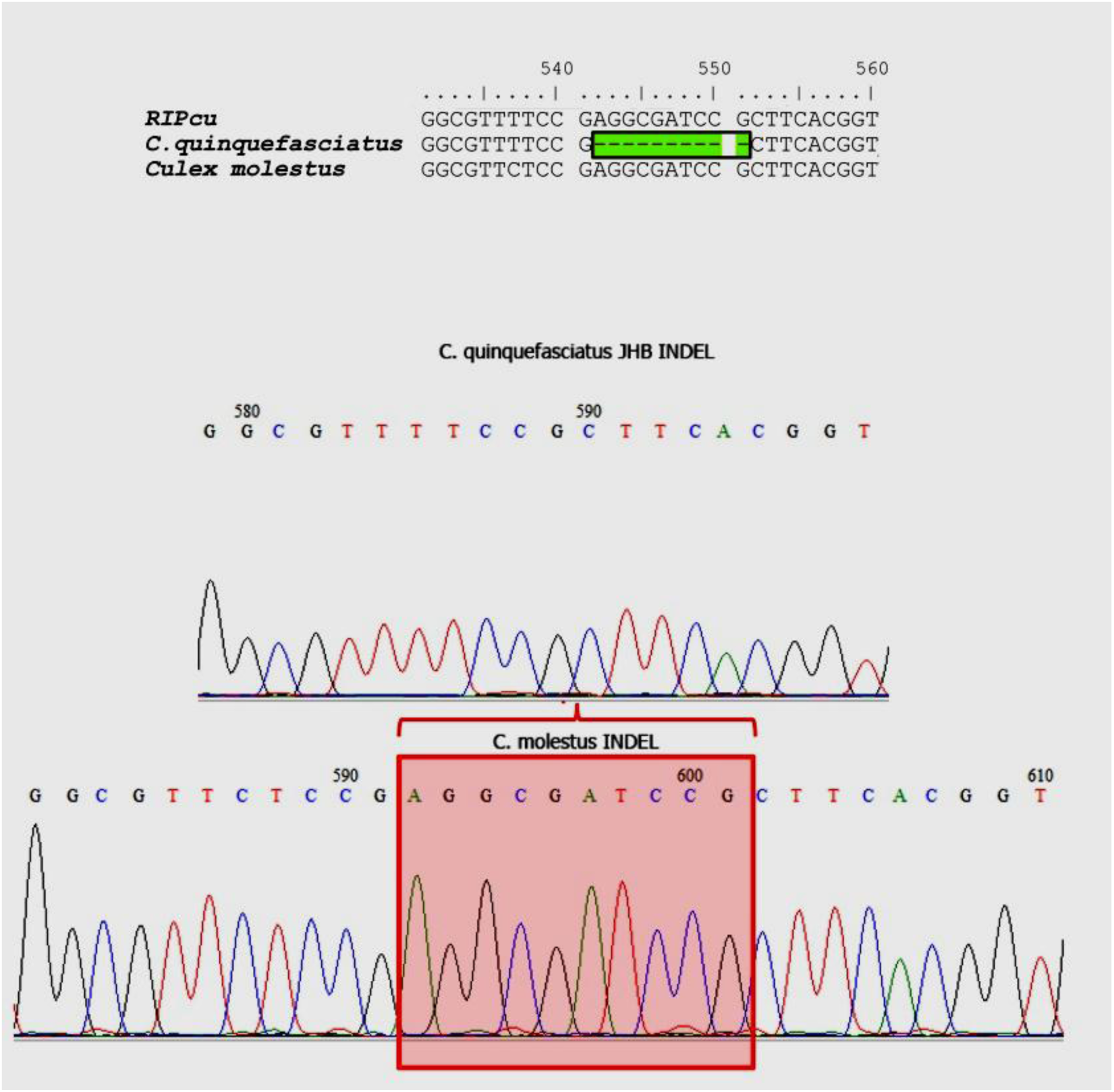
Electropherograms of the region including the deletion of ten nucleotides in *C. quinquefasciatus* JHB MR4 colony. In all cases the region was clearly resolved. Top: sequence of *C. quinquefasciatus* JHB MR4 colony displaying the ten nucleotides deletion. Bottom: Chromatogram of *C. molestus* RIP sequence where no deletion is observed.

**S2 Table:** Orthologs genes between *C. quinquefasciatus*, *A. aegypti* and *An. gambiae*. The GenBank and Vector Base codes are indicated for all protein 1 sequences. The code of each protein used in Fig 3 is shown in the last column.

| ***C. quinquefasciatus*** |  | ***A. aegypti*** |  |  | ***A. gambiae*** |  |  |  |
| --- | --- | --- | --- | --- | --- | --- | --- | --- |
| GenBank | Vector Base code | Gen Bank | Vector Base code | e Value | GenBank | Vector Base code | e Value | Code |
| XP_001850870 | CPIJ009208 |  |  |  |  |  |  | 1 |
| XP_001850871 | CPIJ009209 |  |  |  |  |  |  | 2 |
| XP_001850872 | CPIJ009210 |  |  |  |  |  |  | 3 |
| XP_001850873 | CPIJ009211 | XP_001650164 | AAEL005004 | 1 e-93 |  |  |  | **A** |
| XP_001850874 | CPIJ009212 |  |  |  |  |  |  | 4 |
| XP_001850875 | CPIJ009213 | XP_001650165 | AAEL004999 | 5 e-65 |  |  |  | **B** |
| XP_001850876 | CPIJ009214 |  |  |  |  |  |  | 5 |
| XP_001850877 | CPIJ009215 |  |  |  |  |  |  | 6 |
| XP_001850878 | CPIJ009216 |  |  |  |  |  |  | 7 |
| XP_001850879 | CPIJ009217 | XP_001650161 | AAEL005011 | 0 | XP_309724 | AGAP010970 | 0 | **C** |
| XP_001850880 | CPIJ009218 |  |  |  |  |  |  | 8 |
| XP_001850881 | CPIJ009219 |  |  |  |  |  |  | 9 |
| XP_001850882 | CPIJ009220 |  |  |  |  |  |  | 10 |
| XP_001850884 | CPIJ009222 | XP_001650166 | AAEL005012 | 0 | XP_310862 | AGAP000256 | 0 | **D** |
| XP_001850885 | CPIJ009223 | XP_001650167 | AAEL005003 | 0 | XP_318236 | AGAP010327 | 0 | **E** |
| XP_001850886 | CPIJ009224 | XP_001650168 | AAEL005001 | 6 e-138 | XP_554502 | AGAP010326 | 6 e-87 | **F** |
| XP_001850887 | CPIJ009225 | XP_001650169/70/71 | AAEL005008 | 1 e-153 | XP_318238 | AGAP010325 | 8 e-138 | **G** |
| XP_001850888 | CPIJ009226 | XP_001650173 | AAEL005009 | 0 | XP_318239 | AGAP010324 | 0 | **H** |
| XP_001850889 | CPIJ009227 |  |  |  |  |  |  | 11 |

**S3 Table:** Integrative analyses of substitution rates by SLAC, FEL and REL tests. Those codons 5 under non-neutral selection for each test are presented in violet color. In the consensus column, those 6 codons indicated with blue and dash white inside the box display dN > dS have significant difference. 7 These results were estimated using a p-value < 0.01 for SLAC and FEL tests, and a Bayes factor 8 cutoff = 50 for REL test. Active site codons are indicated in yellow.

| Codon | SLAC dN-dS | SLAC p-value | FEL dN-dS | FEL p-value | REL dN-dS | REL Bayes Factor | Consensus | Codon | SLAC dN-dS | SLAC p-value | FEL dN-dS | FEL p-value | REL dN-dS | REL Bayes Factor | Consensus |
| --- | --- | --- | --- | --- | --- | --- | --- | --- | --- | --- | --- | --- | --- | --- | --- |
| 1 | 0.000 | 0.000 | 0.000 | 1.000 | -0.930 | 3.542.440 | - | 194 | -1.022 | 0.100 | -1.883 | 0.001 | -1.346 | 146.140 | - |
| 9 | -2.072 | 0.043 | -2.006 | 0.001 | -1.461 | 84.762 | - | 207 | -1.499 | 0.004 | -0.877 | 0.000 | -0.957 | 11.951.900 | - |
| 23 | -1.815 | 0.005 | -1.468 | 0.000 | -1.114 | 7.982.030 | - | 213 | -1.081 | 0.090 | -2.813 | 0.002 | -1.219 | 82.171 | - |
| 30 | -1.216 | 0.039 | -483.270 | 0.000 | -1.102 | 477.019 | - | 215 | -1.585 | 0.019 | -0.686 | 0.007 | -0.906 | 341.278 | - |
| 33 | -0.605 | 0.165 | -0.172 | 0.054 | -0.836 | 703.730 | - | 221 | -2.495 | 0.007 | -2.569 | 0.001 | -1.455 | 251.986 | - |
| 34 | -0.405 | 0.247 | -0.120 | 0.078 | -0.764 | 219.770 | - | 222 | -1.783 | 0.023 | -2.166 | 0.002 | -1.559 | 216.213 | - |
| 37 | -1.335 | 0.007 | -0.806 | 0.000 | -0.920 | 2.974.950 | - | 225 | -1.199 | 0.012 | -0.414 | 0.001 | -0.883 | 1.765.240 | - |
| 46 | -1.557 | 0.027 | -0.890 | 0.007 | -0.856 | 86.225 | - | 228 | -1.825 | 0.013 | -2.446 | 0.001 | -2.118 | 433.485 | - |
| 48 | -1.517 | 0.052 | -1.237 | 0.007 | -0.941 | 59.976 | - | 233 | -2.131 | 0.037 | -0.972 | 0.000 | -0.989 | 829.644 | - |
| 49 | -1.485 | 0.012 | -1.696 | 0.000 | -1.178 | 1.589.890 | - | 239 | -1.237 | 0.051 | -2.492 | 0.001 | -1.833 | 672.254 | - |
| 53 | -1.199 | 0.012 | -0.408 | 0.002 | -0.865 | 473.736 | - | 243 | 0.000 | 0.000 | 0.000 | 1.000 | -0.917 | 228.045 | - |
| 55 | -0.902 | 0.112 | -0.546 | 0.026 | -0.895 | 162.204 | - | 244 | -1.468 | 0.013 | -3.293 | 0.000 | -1.342 | 1.222.570 | - |
| 59 | -0.900 | 0.037 | -0.439 | 0.003 | -0.867 | 906.730 | - | 246 | -1.476 | 0.012 | -1.274 | 0.001 | -1.101 | 769.780 | - |
| 61 | -0.300 | 0.333 | -0.073 | 0.121 | -0.575 | 105.687 | - | 247 | -1.199 | 0.017 | -0.808 | 0.000 | -0.970 | 2.253.410 | - |
| 62 | -1.401 | 0.031 | -1.330 | 0.003 | -1.028 | 393.040 | - | 269 | -1.420 | 0.111 | -0.766 | 0.001 | -0.933 | 966.777 | - |
| 66 | -0.502 | 0.215 | -0.256 | 0.039 | -0.848 | 500.612 | - | 274 | -1.324 | 0.019 | -0.790 | 0.003 | -0.914 | 616.562 | - |
| 68 | -1.549 | 0.013 | -1.407 | 0.001 | -1.089 | 652.738 | - | 288 | -0.508 | 0.213 | -0.195 | 0.049 | -0.820 | 380.593 | - |
| 79 | -1.153 | 0.015 | -0.526 | 0.001 | -0.884 | 775.776 | - | 295 | -0.903 | 0.043 | -0.325 | 0.007 | -0.837 | 52.799 | - |
| 82 | -1.787 | 0.005 | -2.016 | 0.000 | -1.146 | 4.806.020 | - | 299 | -1.091 | 0.100 | -0.948 | 0.006 | -0.938 | 183.184 | - |
| 84 | -0.605 | 0.198 | -0.172 | 0.068 | -0.827 | 147.173 | - | 313 | -1.449 | 0.006 | -0.698 | 0.000 | -0.911 | 2.645.140 | - |
| 85 | -1.134 | 0.018 | -0.442 | 0.002 | -0.890 | 945.605 | - | 315 | -1.072 | 0.041 | -0.307 | 0.009 | -0.863 | 777.034 | - |
| 87 | -0.900 | 0.042 | -0.240 | 0.009 | -0.816 | 224.194 | - | 316 | -1.519 | 0.007 | -1.172 | 0.000 | -1.118 | 8.088.370 | - |
| 91 | -1.258 | 0.047 | -0.836 | 0.004 | -0.913 | 255.985 | - | 322 | -1.291 | 0.116 | -2.786 | 0.007 | -1.338 | 24.264 | - |
| 97 | -1.688 | 0.016 | -3.395 | 0.000 | -1.357 | 951.017 | - | 345 | -1.846 | 0.034 | -1.601 | 0.009 | -0.972 | 31.907 | - |
| 120 | -1.170 | 0.076 | -1.137 | 0.007 | -0.928 | 84.017 | - | 365 | -0.564 | 0.177 | -0.131 | 0.072 | -0.763 | 213.515 | - |
| 131 | -0.984 | 0.110 | -5.106 | 0.000 | -2.881 | 528.198 | - | 368 | -1.199 | 0.014 | -0.569 | 0.001 | -0.883 | 1.708.740 | - |
| 135 | -1.207 | 0.038 | -0.600 | 0.008 | -0.871 | 268.034 | - | 370 | -0.581 | 0.200 | -0.443 | 0.004 | -0.854 | 99.841 | - |
| 137 | -1.707 | 0.009 | -0.654 | 0.001 | -0.918 | 976.519 | - | 372 | -1.538 | 0.004 | -0.782 | 0.000 | -0.916 | 1.403.450 | - |
| 140 | -1.628 | 0.004 | -0.589 | 0.000 | -0.907 | 2.990.570 | - | 374 | -1.234 | 0.041 | -0.865 | 0.004 | -0.874 | 214.831 | - |
| 151 | -1.782 | 0.026 | -1.153 | 0.006 | -0.900 | 43.879 | - | 378 | -0.915 | 0.073 | -0.785 | 0.007 | -0.882 | 146.391 | - |
| 153 | -1.377 | 0.033 | -2.082 | 0.001 | -1.264 | 727.823 | - | 380 | -1.012 | 0.050 | -0.434 | 0.014 | -0.819 | 113.302 | - |
| 155 | -1.354 | 0.033 | -1.182 | 0.002 | -0.998 | 214.374 | - | 401 | -0.900 | 0.037 | -0.309 | 0.004 | -0.851 | 588.744 | - |
| 157 | -1.451 | 0.020 | -0.945 | 0.001 | -0.948 | 575.293 | - | 403 | -0.974 | 0.126 | -2.015 | 0.003 | -1.224 | 70.505 | - |
| 158 | -1.065 | 0.025 | -0.661 | 0.001 | -0.890 | 555.446 | - | 412 | -1.138 | 0.018 | -0.436 | 0.002 | -0.897 | 1.059.120 | - |
| 161 | -0.816 | 0.050 | -0.316 | 0.007 | -0.839 | 219.591 | - | 415 | -1.011 | 0.096 | -0.321 | 0.058 | -0.838 | 120.470 | - |
| 165 | -1.676 | 0.006 | -1.468 | 0.000 | -1.211 | 2.108.320 | - | 419 | -2.660 | 0.032 | -0.662 | 0.004 | -0.927 | 770.996 | - |
| 166 | -0.585 | 0.171 | -0.198 | 0.046 | -0.816 | 400.256 | - | 429 | -2.170 | 0.007 | -4.440 | 0.000 | -1.589 | 950.750 | - |
| 169 | -0.502 | 0.215 | -0.204 | 0.049 | -0.839 | 429.773 | - | 430 | -1.078 | 0.038 | -0.277 | 0.011 | -0.846 | 536.350 | - |
| 175 | -1.604 | 0.070 | -1.654.870 | 0.001 | -3.108 | 191.650 | - | 431 | -1.105 | 0.110 | -11.486 | 0.002 | -1.226 | 24.637 | - |
| 178 | -0.786 | 0.117 | -0.678 | 0.003 | -0.888 | 73.938 | - | 435 | -0.839 | 0.087 | -1.752 | 0.007 | -1.001 | 89.806 | - |
| 180 | -1.199 | 0.012 | -0.602 | 0.000 | -0.901 | 4.823.560 | - | 436 | -1.637 | 0.011 | -0.804 | 0.001 | -0.973 | 1.717.340 | - |
| 181 | -0.557 | 0.179 | -0.149 | 0.064 | -0.791 | 388.112 | - | 443 | -0.871 | 0.115 | -0.943 | 0.010 | -0.873 | 52.265 | - |
| 185 | -1.757 | 0.005 | -2.482 | 0.000 | -1.103 | 6.544.390 | - | 458 | -1.344 | 0.033 | -1.325 | 0.001 | -1.084 | 464.844 | - |
| 187 | -1.075 | 0.039 | -0.313 | 0.009 | -0.848 | 573.723 | - | 473 | -1.271 | 0.048 | -1.339 | 0.004 | -0.976 | 129.196 | - |
| 188 | -1.199 | 0.012 | -0.615 | 0.000 | -0.914 | 2.766.090 | - | 499 | -0.602 | 0.195 | -0.154 | 0.065 | -0.790 | 250.543 | - |
| 190 | -1.419 | 0.006 | -0.777 | 0.000 | -0.928 | 8.231.280 | - | 500 | -2.521 | 0.001 | -3.285 | 0.000 | -1.580 | 10.402.000 | - |
| 191 | -1.178 | 0.029 | -0.417 | 0.005 | -0.880 | 1.466.510 | - |  |  |  |  |  |  |  |  |
| 192 | -1.049 | 0.045 | -0.702 | 0.003 | -0.880 | 546.106 | - |  |  |  |  |  |  |  |  |
| 193 | -1.349 | 0.018 | -0.798 | 0.002 | -0.871 | 349.318 | - |  |  |  |  |  |  |  |  |

